# The 2D and 3D ultrastructure of symbiosomes and associated vesicular structures in *Lotus japonicus* root nodule symbiosis

**DOI:** 10.64898/2026.05.03.722514

**Authors:** Isabella Gantner, Katarzyna Parys, Andreas Klingl

## Abstract

In root nodule symbiosis, symbiosome compartments accommodate nitrogen-fixing rhizobia inside the plant cell. Differentiated into bacteroids, the rhizobia are surrounded by a peribacteroid space and a plant-derived peribacteroid membrane, which separates them from the plant cytoplasm but allows signal and nutrient exchange between host and microbe. The morphological features of symbiosomes are primarily determined by ultrastructural single focal plane imaging, with limited information about spatial details. This study combines 2D and 3D imaging, using transmission electron microscopy and focused ion beam scanning electron microscopy as complementary techniques to analyse the symbiosome ultrastructure and organisation in *Lotus japonicus* wild-type plants. The 3D model of a mature colonised root nodule cell region demonstrates a dense, puzzle-like arrangement of symbiosomes relative to one another and adjacent plant organelles. The symbiosome shape and size depends on the orientation and number of bacteroids within the compartment and features connective tubular structures. Furthermore, vesicular structures, some likely of bacterial origin, were present at the interface. The study presents a multi-angled analysis of symbiosome-related structures, highlighting their volumes, spatial distribution, and pronounced compactness. Interface associated vesicles, protrusions and connective structures hint towards a dynamic and flexible system that contributes to the plant-microbe crosstalk.

## Introduction

In root nodule symbiosis, rhizobia are accommodated within living plant cells, providing mutual benefits to both the host and microsymbiont (Brewin, 1991). By reducing atmospheric nitrogen to ammonia (Bergersen, 1965), the bacteria satisfy the nitrogen needs of the plant, which is essential for its growth and development (Kondorosi *et al*., 2013). In return, the plant transports dicarboxylic acids such as malate, fumarate, or succinate to the microbe and represents an attractive carbon source for the inhabitants (Bapaume & Reinhardt, 2012).

A hallmark of this symbiosis is the nodule, a root organ formed *de novo* that hosts rhizobia and provides an oxygen-depleted environment for nitrogen fixation. Legumes develop either indeterminate or determinate nodules, depending on the host species. Indeterminate nodules are found in *Medicago, Pisum* and *Vicia*, and are characterised by a persistent apical meristem and distinct developmental zones within the nodule (Vasse *et al*., 1990). In contrast, determinate nodules are formed by *Glycine* and *Lotus* and lack an apical meristem. Instead, the nodule contains mature, enlarged, nitrogen-fixing symbiotic cells at a similar developmental stage (Kondorosi *et al*., 2013). In both nodule types, mature symbiotic cells can accommodate a high number of rhizobia to enhance nitrogen-fixation efficiency. This requires major changes in the spatial organisation of cellular compartments, which may be induced by a distinct reticulate array of cytoplasmic actin that develops after infection (Whitehead *et al*., 1998). Furthermore, the host cytoplasm contains numerous mitochondria, dictyosomes and endoplasmic reticulum (ER) (Kijne & Pluvqué, 1979), reflecting high metabolic activity of the nodule cell, while the large central vacuole breaks into smaller fragments to create space for accommodation of the rhizobia (Bapaume & Reinhardt, 2012; Gravin *et al*., 2014).

Rhizobia are taken up by the host plant through a root hair infection thread, which is a tubular structure that guides the bacteria through the root epidermis and towards cortical cells of a newly developing nodule (Spronsen *et al*., 2001). In the target cells of the central nodule tissue, the rhizobia are released by infection droplets which bud off the thread (Tsyganova *et al*., 2021). This results in an endocytotic uptake of the rhizobia into a cytoplasmic compartment, the symbiosome, where a plasma membrane-derived peribacteroid membrane (PBM) encloses the rhizobia (Pankhurst *et al*., 1979). Inside this organelle-like compartment, the bacteria undergo growth, cell division and differentiation into bacteroids. The physiology, morphology, and number of bacteroids within a symbiosome are highly diverse and depend on the plant host (Dart & Mercer, 1966). For example, *Sinorhizobium meliloti*, the endosymbiont of *Medicago truncatula*, forms elongated, swollen or spherical bacteroids, whereas the bacteroid morphology of *Mesorhizobium loti* that nodulates *Lotus japonicus* is unaffected compared to their free-living form (Mergaert *et al*., 2006). In a symbiosome, bacteroids are surrounded by a peribacteroid space (PBS), which together with the PBM forms the plant-microbe interface (PMI), a zone in direct contact with both the plant cytoplasm and the bacteroid outer membrane (Bergersen & Briggs, 1958). The PMI acts as a central hub for crosstalk and nutrient exchange between the interaction partners. The PBS is filled with diffuse, granular material (Bergersen & Briggs, 1958) and contains a variety of compounds, including enzymes (Mellor *et al*., 1984), proteins (Katinakis *et al*., 1988), and sugars (Tejima *et al*., 2003), derived from both the plant and the bacteroid. Additionally, extracellular vesicles are often located in the space (Ayala-Garcia *et al*., 2025), which, besides transporters and channels located in the PBM (Jeong *et al*., 2004; Clarke *et al*., 2014) may be involved in substance exchange and communication between the interaction partners through the interface (Kijne & Pluvqué, 1979).

The detailed description of the symbiosome interface and the morphology of intracellular accommodation structures is heavily dependent on microscopy techniques (Reagan *et al*., 2018). The first morphological descriptions of bacteroids in *Vicia* (Becker, 2017) can be traced to the pioneering article by Martinus Willem Beijerinck published in 1888 (Beijerinck, 1888),. Later, Bergersen and Briggs published in 1958 one of the first descriptions of the 2D ultrastructure and organisation of symbiosomes in *Glycine* root nodules cells (Bergersen & Briggs, 1958). Since then, several studies focused on intracellular root nodule structures primarily in the genera *Medicago, Lotus* and *Glycine* using light, fluorescence, and transmission electron microscopy (TEM) combined with two-dimensional (2D) image analysis. However, in the last few decades, the field of structural biology has been revolutionised by advancements in microscopy. Techniques such as Serial Block-Face Scanning Electron Microscopy and Focused Ion Beam Scanning Electron Microscopy (FIB/SEM) have facilitated the creation of high-resolution image stacks. These three-dimensional (3D) volume datasets provide a detailed, multi-angled view of specific structures, allowing researchers to answer complex questions, such as the spatial distribution or connectivity between structures that are impossible to address with 2D image data. One of the first applications of FIB/SEM in the biological sciences was demonstrated in 1993 by Young and his colleagues, who employed the technique to characterise the internal structures of arthropods (Young *et al*., 1993; Reagan *et al*., 2018). Since then, FIB/SEM has been widely adopted as a powerful tool for elucidating the ultrastructural details of several biological tissues in human and plant research (Schmidt *et al*., 2011; Crumpton-Taylor *et al*., 2012).

Strikingly, despite the availability of advanced spatial imaging techniques, there is limited 3D ultrastructural data on rhizobia-legume root nodule symbiosis (Nakhforoosh *et al*., 2024). There were initial attempts to visualise infected root nodule cells in 3D using SBF/SEM (Sánchez-Cañizares *et al*., 2026) and FIB/SEM (Reagan *et al*., 2018). However, the ultrastructure of symbiosomes, bacteroid organisation, and the PBS was not described in detail. This study elucidates the ultrastructural spatial arrangement of bacteroids and symbiosomes in the tissue of *L. japonicus* nodules colonised by *M. loti*. By combining 2D and 3D data, we reveal the complex structural arrangement of symbiosomes and plant organelles within a symbiotic cell and explore the symbiosome shape, number, and volume. Furthermore, this study provides a detailed overview of the locations of symbiosome-associated vesicles and their origin using TEM and immunogold labelling.

## Materials and Methods

### Plant growth conditions and rhizobium inoculation

Seeds of *L. japonicus* ecotype Gifu B-129 wild-type (Handberg & Stougaard, 1992) were scarified and surface-sterilised for 5 min in a solution containing 2 % NaClO and 0.01 % (w/v) SDS. After five washes with sterile water, seeds were incubated overnight at 4 °C. They were then plated on 0.8 % water-agar square plates, sealed with microtape and kept in the dark for two days before transferring to a PolyClima growth cabinet set to 24 °C and 50 % humidity under long day conditions (16 h light, 8 h dark; 160 μmol s^-1^ m^-2^).

Seven-day-old seedlings were inoculated with a suspension of *M. loti* MAFF 303099 *Ds*Red, adjusted to a final optical density at 600 nm of 0.05 in FABnod medium with reduced nitrogen content (500 µM MgSO_4_·7H_2_O, 250 µM KH_2_PO_4_, 250 µM KCl, 250 µM CaCl_2_·2H_2_O, 100 µM KNO_3_, 25 µM Fe-EDDHA, 50 µM H_3_BO_3_, 25 µM MnSO_4_·H_2_O, 10 µM ZnSO_4_·7H_2_O, 0.5 µM Na_2_MoO_4_·2H_2_O, 0.2 µM CuSO_4_·5H_2_O, 0.2 µM CoCl_2_·6H_2_O; pH 5.7). For inoculation, the root of each seedling was dipped into the bacterial suspension and then transferred to square plates containing solid FABnod medium with 0.8 % agar. Plates were sealed with micropore tape, and roots were covered with black foil to prevent light exposure. The plates were transferred back to the PolyClima growth cabinet and placed in a tilted position. Root nodules were harvested after 14 days post-inoculation (dpi) and incubated in fixation solution according to the respective preparation protocol.

### Transmission electron microscopy

Sample preparation for TEM was carried out by fixing 14 dpi *L. japonicus* root nodules with 2.5 % (w/v) glutaraldehyde in 75 mM cacodylate buffer containing 2 mM MgCl_2_ (Espinoza-Corral *et al*., 2019) for 4 days at 4 °C. Chemical fixation was used as it already delivered excellent results in previous studies and it should be ensured that the results are fully comparable. The plant material was washed with fixation buffer and distilled water three times each at room temperature before continuing with a post-fixation step with 1% osmium tetroxide (w/v) for 2 h. In the next step, dehydration in a graded acetone series from 10-100 %, including additional contrasting with 1 % uranyl acetate (UrAc) (w/v) for 20 min in the 20 % (v/v) acetone step, was performed. The samples were gradually infiltrated with Epon resin 812 and kept in the 100 % resin step overnight. Final embedding was performed in flat embedding molds, and samples were polymerised overnight at 63 °C. With an ultracut machine (Reichert) and a diamond knife (DiATOME), the polymerised samples were sectioned in 50-70 nm sections, mounted on collodion-coated Cu grids and post-stained with 80 mM lead citrate (pH 13). Images were generated using a Zeiss EM912 (Carl Zeiss Microscopy GmbH), operated at 80 kV, and equipped with a Tröndle 2 k × 2 k slow-scan CCD camera (TRS, Tröndle Restlichtverstärkersysteme, Moorenweis, Germany).

### Immunogold labelling

Root nodules of *L. japonicus* were prepared for TEM as described above, with the exception that 4 % paraformaldehyde was used for fixation. The less sensitive Epon resin 812 was chosen for this experiment due to its excellent structural preservation, which is particularly important for the labelling of small structures such as vesicles and their membranes (Flechsler *et al*., 2020). Freshly sliced thin sections of the embedded nodule were placed on nickel grids and incubated in a drop of 0.1 % Glycin in phosphate buffer saline (PBS) (137 mM NaCl, 2.7 mM KCL, 8.4 mM Na_2_HPO_4_, 1.4 mM KH_2_PO_4_; pH 7.4) and a drop of 1 % bovine serum albumin (BSA) in PBS for 5 min each. The Lipid A primary antibody (Thermo Fisher; PA1-73178) was diluted 1:1000 in 0.1 % BSA in PBS. Samples were incubated with the antibody for 1.5 h and washed four times with 0.1 % BSA in PBS. The samples were then incubated for 1 h with the secondary antibody (anti-goat raised in rabbit, coupled to colloidal gold particle of 10 nm, Sigma) diluted 1:20 in 0.1 % BSA in PBS for 1 h. Afterwards, the samples were washed five times in 0.1 % BSA in PBS, twice in PBS and thrice in sterile water for 2 min each. To improve contrast, the samples were stained with 2 % UrAc for 15 min and washed twice with sterile water. Between all washing, antibody and staining steps, grid blotting on filter paper was repeated to remove the liquids. To assess the specificity of the secondary antibody, a control was processed with the same protocol but without the primary antibody step. Images of the immunogold labelled samples and the control were taken with the TEM. The experimental procedure was repeated three times independently.

### Negative staining

Gold grids were coated with a carbon film of 10 nm and additionally hydrophilized. The grids were transferred to an Erlenmeyer flask containing 50 ml of TY-CaCl_2_ media with Gentamycin [12.5 μg/mL] and inoculated with *M. loti* MAFF 303099 *Ds*Red. The bacterial culture was incubated at 28 °C and 180 rpm. After two days, grids overgrown with bacteria were removed, blotted on filter paper and prepared for negative staining. For staining, the carbon-coated side of the grids was washed twice with sterile water, then stained for 10 s with 0.5 % UrAc. Between all washing and staining steps, blotting was repeated to remove the liquids. Images of the negatively stained samples were taken with TEM.

### Focused ion beam electron microscopy

Samples for FIB/SEM series were pretreated, fixed, and embedded as described for TEM. After polymerisation for 96 h at 63 °C, the trimmed resin block was mounted on a thin aluminum pin and coated with 10 nm carbon. Milling and imaging of the samples were carried out with an Auriga 40 Crossbeam workstation using the SmartSEM software package (Carl Zeiss Microscopy GmbH). Before milling the sample region of interest was coated with a platinum protection deposition for 14 min at 2 nA. Images were recorded at an accelerated voltage of 1.5 kV using an energy-selective backscattered electron detector (EsB) with the EsB-grid set to 500 V and the 30 μm aperture. The scan speed was set to an exposure time of 80 s per image with a total resolution of 2.048 × 1.536 pixels. An ion beam current of 100 pA was used for milling with 2 nm of material being milled away in each cycle. Images were always acquired in multiples of 2 nm. The voxel size range was 8 nm in x/y and 8 nm in z.

### Segmentation and 3D reconstruction

The 3D reconstruction of a colonised *L. japonicus* nodule cell region was performed using the image analysis software Amira (Amira 3D 2024.1; ThermoFisher Scientific). Prior to 3D visualisation, the 460 TIF images were transformed and aligned into a single TIF file using the open-source image processing platform FIJI (ImageJ 1.53e). In Amira, each plant organelle, including symbiosome membranes and bacteroids, was drawn as a single material in the segmentation editor with the help of an external creative pen display (Wacom Cintiq 22; Wacom K.K., Kazo, Saitama, Japan). Visualisation of the materials was conducted using the application tools: Generate surface and Surface view. To display the structures in their respective Surface views, the drawing style for the bacteroid and vesicle material was set to shaded, except for the symbiosome membranes, which were set to transparent. Volume calculations of the individual structures were conducted using the application tool Material statistics.

### 2D data analysis

TEM images were acquired of the outer colonised tissue (three cell layers) of three root nodules, each from a different plant (sample preparation and image generation as described above). Images of ten cells per outer region were taken with a constant image frame dimension. Only symbiosomes that were completely contained within the image frame (10.4 μm x 10.4 μm) were included in the measurements. The area and number of bacteroids and symbiosomes were quantified in FIJI (ImageJ 1.53e) in combination with an external creative pen display (Wacom Cintiq 22; Wacom K.K., Kazo, Saitama, Japan) to trace the outlines of the structures of interest. To compare inner and outer root nodule cell regions, the same analysis was performed for cells located in the centre of one of the root nodule samples.

### Statistics

Graphs and statistical analyses were generated with the R Studio software (Version 1.1.463). Data was converted into numeric values to ensure compatibility with statistical analyses. The dataset was converted into long format and distribution was assessed using the Shapiro–Wilk test for normality, while variance homogeneity was tested with an F-test. A standard two-sample t-test or non-parametric Mann–Whitney U test was applied to compare measurements between groups. Statistical significance was evaluated at a level of p ≤ 0.05.

## Results

### *Mesorhizobium loti* in its free-living state and accommodated in symbiosomes

The morphology of *M. loti* in its free-living state was analysed using negative staining and TEM. The cultured cells showed a rod shape of 1-1.5 µm in length (Fig. 1a) and a single flagellum (length 6-7 µm) as well as lateral pili (length 450 nm) extruding from the cell (Fig. 1b). The morphology of free-living cells was compared to differentiated *M. loti* bacteroids by an ultrastructural analysis on cross-sections of mature *L. japonicus* root nodules (14 days post-inoculation with *M. loti)* (Fig. 1c-d). The nodules were selected based on their diameter (1-2 mm) and pink colouration, indicating leghaemoglobin production (Bergersen & Goodchild, 1973). TEM micrographs of nodule cross-sections revealed the presence of numerous bacteroids encapsulated within symbiosome compartments separating the microbes from the plant cytoplasm. Every symbiosome was engulfed by an electron-dense PBM surrounding one or multiple bacteroids, surrounded by an electron-translucent PBS (Fig. 1e).

**Fig. 1.**
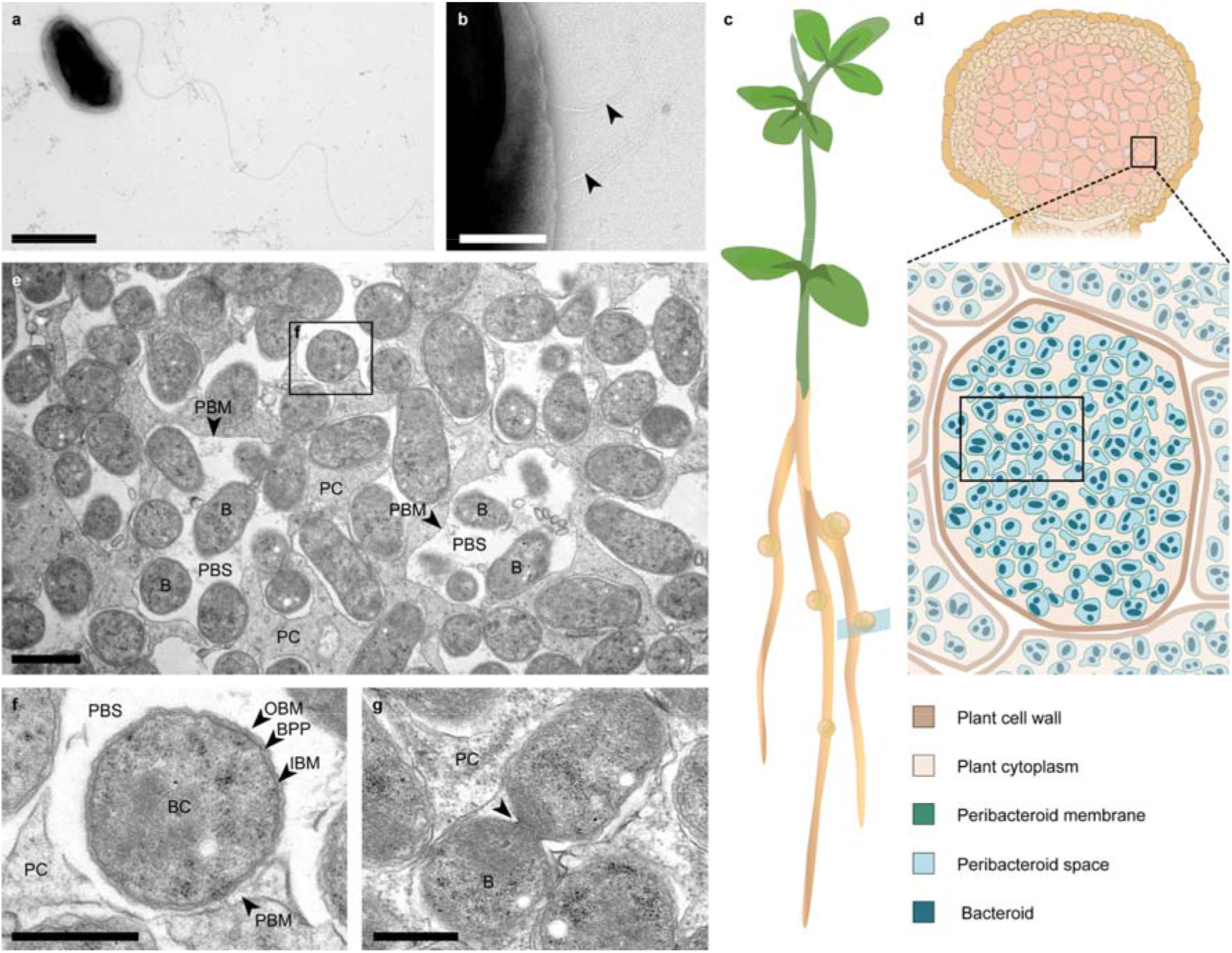
Ultrastructural characteristics of *Mesorhizobium loti* and its accommodation in *Lotus japonicus* root nodule cells. **a)** *M. loti* cell in a free-living state and with a single flagellum (scale bar: 1 µm) and **b)** a *M. loti* cell with several pili on the lateral side of the cell, visualised by negative staining and TEM (scale bar: 200 nm). **c)** Schematic drawing of an *L. japonicus* plant with several root nodules and **d)** a cross-section of a colonised determinate root nodule. Cells in the central tissue colonised by *M. loti* are highlighted in pink, with a close-up view of a cell of the outer colonised tissue layer, densely packed with symbiosomes. The black frame in the close-up symbolises the tissue region where image **e)** and also the FIB/SEM stack for 3D reconstruction was generated. **e)** TEM image of *L. japonicus* root nodule cell region in the outer cell layer of the nodule tissue. The region is densely colonised by bacteroids enclosed in symbiosomes, which comprise a plant-derived PBM, the PBS and one or multiple bacteroids (scale bar: 1 µm). **f)** Close-up cross-section of a single bacteroid with its inner and outer bacteroid membranes as well as the bacteroid periplasm. The bacteroid is surrounded by PBS (scale bar: 500 nm). **g)** TEM image of a longitudinal section of a bacteroid undergoing cell division (scale bar: 500 nm). Labelling: B: bacteroid; BC: bacteroid cytoplasm; BPP: bacteroid periplasm; IBM: inner bacteroid membrane; OBM: outer bacteroid membrane; PBM: peribacteroid membrane; PBS: peribacteroid space; PC: plant cytoplasm.

The close-up cross-section of a symbiosome visualises that three membranes (PBM-OBM-IBM) were present as well as the PBS between the cytoplasm of the plant host and the cytoplasm of the gram-negative *M. loti* (Fig. 1f).The overall morphology of *M. loti* did not change after differentiation into a bacteroid compared to its free-living state. Cells maintained their rod shape, and cell division events were observed (Fig. 1g). However, bacterial cell appendages were absent for cells accommodated in a symbiosome.

In order to evaluate, if the size and amount of symbiosomes and accommodated bacteroids depends on the location of the cell in the *L. japonicus* nodule tissue, a 2D TEM image-based comparison of cells from the inner and outer colonised cell layers was performed (Supplementary Fig.1). The results showed no significant difference for symbiosome-related characteristics dependent on the cell position within the nodule. Determinate nodules lack a persistent meristem and distinct developmental zones, and all infected cortical cells are at the same developmental stage (Kondorosi *et al*., 2013). As this consistency is also evident in the ultrastructural characteristics of the symbiosomes, colonised plant cells of the outer layers of the nodule tissue were used as representative tissue (Fig. 1d) for further 2D and 3D analysis of symbiosomes.

### Tight spatial arrangement of symbiosomes and plant organelles within the nodule cell

Using a FIB/SEM image stack of a colonised *L. japonicus* cell region in the outer cell layer of the infected root nodule tissue (Fig. 2a), a 3D model was generated through structural segmentation (Fig. 2b). The model includes 56 symbiosomes agglomerated in the plant cytoplasm. Each symbiosome was assigned a distinct colour with the PBM rendered in transparent colour and bacteroid cells in opaque colour. The top view (Fig. 2b) and perspective side view (Fig. 2c) of the entire model show the relative spatial arrangement of the symbiosomes. They are tightly packed and arranged in interlocking positions. The perspective view of isolated bacteroids without the surrounding PBM and PBS (Fig. 2d) shows that the microbes are tightly clustered together, efficiently occupying the available space without following a specific repetitive orientation.

**Fig. 2.**
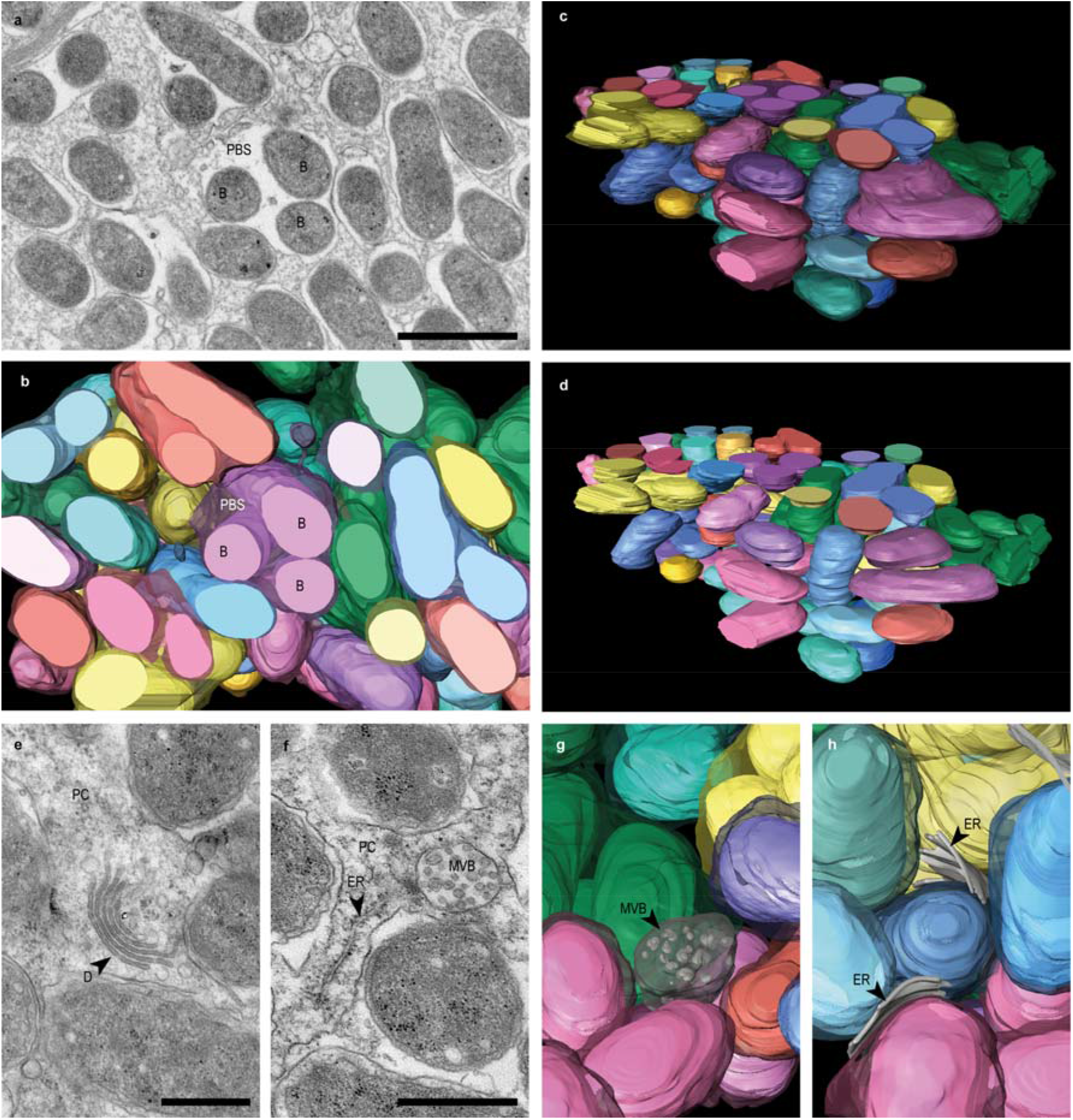
The tight spatial arrangement of symbiosomes and small plant organelles in a colonised root nodule cell. FIB/SEM image stack dataset of a colonised *L. japonicus* cell region in the outer cell layer of the infected root nodule tissue. **a)** A single 2D image of the FIB/SEM stack with several symbiosomes (scale bar: 1 µm) and corresponding 3D model of this colonised region in **b)** a top view, and **c)** a perspective side view. The symbiosome and the bacteroids it contains share the same colour. The PBM is rendered in transparent colours, and bacteroid cells (outer bacteroid membrane) in opaque colours. Symbiosomes contain one or more bacteroids. The model visualises a tight, interlocking arrangement of symbiosomes. **d)** Corresponding rendering of the stack in **c)** showing only the bacteroid spatial arrangement without the PBM. **e-h)** Spatial arrangement of plant cell organelles between symbiosome compartments in the plant cytoplasm visualised in 2D TEM images: **e)** a dictyosome, **f)** rough endoplasmatic reticulum and a multi-vesicular body (scale bars: 500 nm), and in the corresponding 3D model **g-h)**, visualizing the adjustment of small plant organelles placement to the limited space in the remaining plant cytoplasm. Labelling: B: bacteroid; D: dictyosome, ER: endoplasmic-reticulum; MVB: multi-vesicular body; PBS: peribacteroid space; PC: plant cytoplasm.

Plant cytoplasm between symbiosomes contains different organelles that support the cell metabolic activity of the host cell and promote symbiosome maintenance (Kijne & Pluvqué, 1979) (Bapaume & Reinhardt, 2012). The thin layer of host cytoplasm between the symbiosomes contains plant organelles like dictyosomes (Fig. 2e), rough ER and multi-vesicular bodies (Fig. 2f), which are visible in both 2D and 3D data. However, only the 3D reconstruction depicts the confinement of these small plant organelles between the symbiosomes (Fig. 2g-h) and visualizes, that structures like the rough ER are tightly packed between the symbiosome compartments (Fig. 2h).

### Symbiosome shape variations and volumes

FIB/SEM stack analysis provides an unprecedented level of spatial information on mature root nodule cells filled with symbiosomes. Capitalising on this, the structure, shape and size of individual symbiosomes within the 3D model were analysed. A symbiosome consists of at least one rod-shaped *M. loti* bacteroid surrounded by a thin layer of PBS and a PBM which follows the shape of the bacteroid in close distance (Fig. 3a). Because of the bacteroid shape imitation by the PBM, the shape variety of symbiosomes depends on the bacteroid arrangement inside the compartment. It can range from small symbiosomes with only one bacteroid, to larger pouch-like symbiosomes with multiple centrally clustered bacteroids (Fig. 3b) or to symbiosomes with chain-like distribution (Fig. 3c), where several bacteroids each have their own envelope unit, but share the same PBS and PBM because of tubular connections (diameter of 50 nm) (Fig. 3d-e). These connections were present for six symbiosomes in the 3D model.

**Fig. 3.**
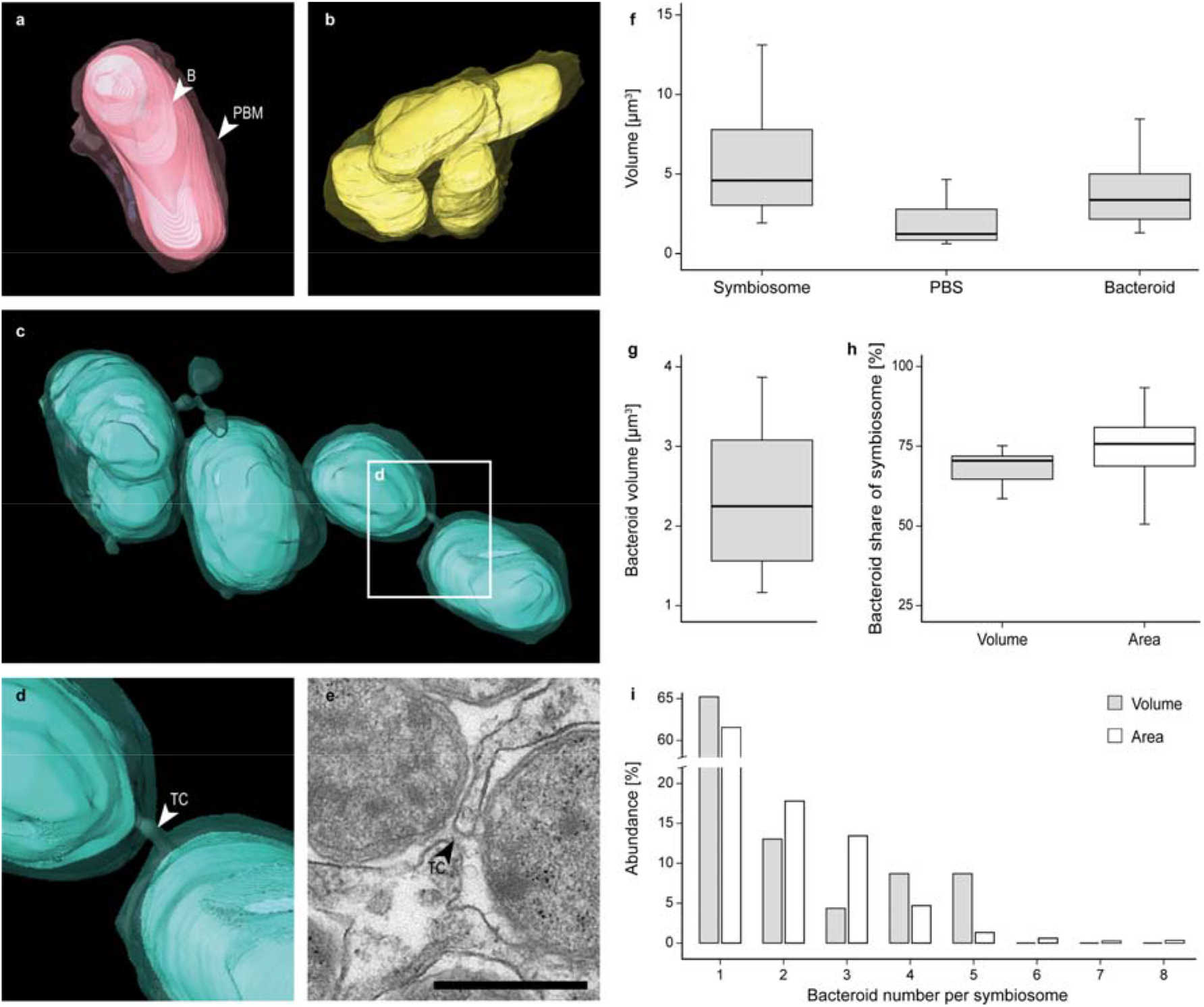
Variation in symbiosome shape, bacteroid number and volume in *Lotus japonicus* root nodules. The rod-shaped *M. loti* bacteroids are surrounded by a thin layer of PBS and a PBM, both tightly following the shape of the bacteroids. Symbiosome shape largely depends on the number and orientation of bacteroids. **a)** Symbiosome with a single bacteroid. **b)** Symbiosome with multiple tightly packed bacteroids. **c)** Symbiosome that contains multiple bacteroids which are spatially organised with respect to each other while sharing the same PBS and PBM because of connective structures. **d)** Close-up of panel **c). e)** A TEM image, visualising the connective structure as a small tube formed by the PBM (scale bar: 500 nm). All symbiosome-related measurements used for the following graphs are based on the data generated from the 3D reconstruction (grey) and from 2D TEM images analysis (white). **f)** The symbiosome (n = 23), PBS and total bacteroid volume, measured from structures which were completely captured in the FIB/SEM stack. **g)** Volume measurements of the individual bacteroid cells (n = 42) and **h)** the bacteroid volume share (n = 23) and bacteroid area share of a symbiosome (n = 1512). **i)** The abundance of symbiosomes with a certain bacteroid number in the 3D (n = 23) and 2D (n = 1512) data. Labelling: B: bacteroid; PBM: peribacteroid membrane; TC: tubular connection.

The volume measurements of 23 complete symbiosomes captured in the FIB/SEM stack demonstrate that a symbiosome compartment has a median volume of 4.6 µm^3^. This is further divided into a PBS volume of 1.2 µm^3^ and a total bacteroid volume of 3.4 µm^3^ (Fig. 3f). For comparison, the volume of a single *M. loti* bacteroid cell had a volume of 2.2 µm^3^ (Fig. 3g).

To compare the structural information obtained from 3D reconstructions with 2D approaches, an additional 2D TEM image-based dataset was acquired from 10 cells each of three independent nodules (n = 3). These datasets further served to evaluate whether symbiosome structural proportions derived from 3D analysis of a single cell region are conserved across multiple cells within the same tissue layer.

The number and area of symbiosomes (n = 1512) and their enclosed bacteroids (n = 2584) were measured, and structural proportions were compared to the 3D volume data. The area proportions for symbiosome related structures (Supplementary Fig. 2) showed a similar distribution for PBS and total bacteroid area, as it was observed for the volume data (Fig. 3f). Both approaches showed that the bacteroids occupy the largest volume (in 3D data analysis) and area (in 2 D data analysis) share of the total symbiosome compartment, accounting for 70.5 % of the compartment’s volume and 75.7 % of its area (Fig. 3h). The 2D dataset acquired from symbiosome sections with variable cutting axes, however, showed a higher data distribution then the 3D dataset for the volume share. Furthermore, both datasets show that symbiosomes containing a single bacteroid are the most abundant category in the *L. japonicus* root nodule tissue at this developmental stage (Fig. 3i), with 65.2 % for the 3D dataset and 61.6 % for the 2D dataset. The abundance of symbiosomes including multiple bacteroids decreases with increasing number of bacteroids. Symbiosomes containing more than five bacteroids were rare. The data shows an approximately exponential decay, which is particularly evident in the 2D dataset with restricted morphological information of a single focal plane, but a representative large sample size.

The tendency towards symbiosomes containing a low number of bacteroids results in a high degree of compartmentalisation within the plant cell. Moreover, this pattern reflects a pronounced level of cellular compactness in colonised root nodule cells, as the tight packing occurs not only between symbiosomes but also within them, with densely packed bacteroids occupying most of the internal space.

### Vesicular structures associated with the symbiosome interface

The ultrastructural 2D and 3D analysis of infected nodule cells revealed vesicle-like structures associated with symbiosomes. These spherical compartments were mostly homogeneous in shape, surrounded by a lipid bilayer and appeared to contain electron dense, heterogeneous cargo (Fig. 4). The vesicles were observed as free vesicles in the plant cytoplasm or peribacteroid space, or associated with the bacteroid or peribacteroid membrane. Vesicles associated with the peribacteroid membrane were present on both the plant and microbial sides of the symbiosome interface, thereby expanding the luminal and cytoplasmic membrane compartments (Fig. 4a). As documented in the 3D model and TEM images, vesicles from the plant cytoplasm side of the interface were attached to the PBM surface (Fig. 4b) or partly embedded in the PBM (Fig. 4c), hereafter “external vesicles”. Furthermore, vesicles were also present inside the PBS or encapsulated in a connected multi-vesicular body-like compartment, sharing the same PBS (Fig. 4d), hereafter “internal vesicles”. Internal vesicles were further categorised according to their location: 1) vesicles attached exclusively to the PBM (Fig. 4e), 2) free vesicles in the PBS, 3) vesicles attached simultaneously to the PBM and outer bacteroid membrane (Fig. 4e), and 4) vesicles attached exclusively to the outer bacteroid membrane (Fig. 4f).

**Fig. 4.**
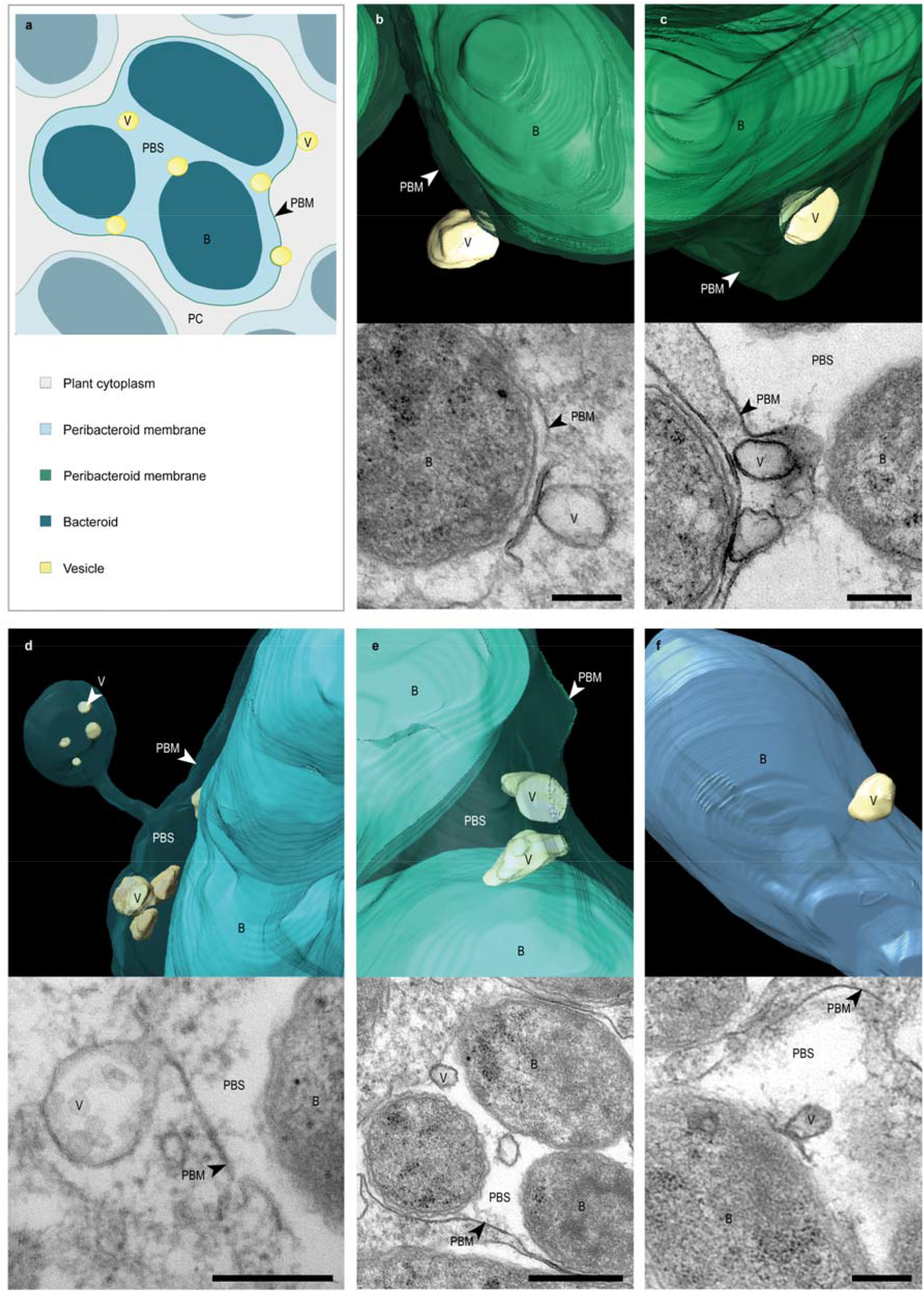
Vesicular structures within the peribacteroid space and attached to the bacteroid- and peribacteroid membrane of symbiosomes. **a)** Schematic drawing summarising the observed vesicle locations associated with symbiosomes. **b-e)** 3D reconstruction of vesicle locations alongside TEM images. **b)** Vesicle attached to the outer surface of the PBM and **c)**, partly surrounded by the PBM (scale bar: 200 nm). **d)** Vesicles located free in the PBS or in a separated compartment with a connection to the main symbiosome corpus (scale bar: 200 nm). **e)** Vesicles in the PBS attached to the inner surface of the PBM or simultaneously to the PBM and outer bacteroid membrane (scale bar: 500 nm). **f)** Vesicle attached to the bacteroid outer membrane (scale bar: 200 nm). Labelling: B: bacteroid; PBM: peribacteroid membrane; PBS: peribacteroid space; V: vesicle.

Of the 23 complete symbiosomes in the 3D model, 13 contained external or internal vesicles. For symbiosomes containing internal vesicles, the median share of vesicle volume in the PBS was 1.6 % (Fig. 5a). There was no significant difference in median volume between internal vesicles (n = 7; median volume of 8.3 × 10^-3^ µm^3^) and external vesicles (n = 30; median volume of 8.13 × 10^-3^ µm^3^) (Fig. 5b). However, the wide spread of values in both categories indicates substantial heterogeneity in vesicle size. With respect to the total vesicle count, internal vesicles (71.4 %) were more abundant than external vesicles (28.6 %) (Fig. 5c-d). Most internal vesicles were attached exclusively to the PBM membrane (60 %) or free in the PBS (24 %) (Fig. 5e), while only a small proportion (16 %) was attached to a bacteroid. The analysis of vesicle abundance in relation to their location in individual symbiosomes shows that in 11 (84.6 %) of the 13 symbiosomes, external vesicles were attached to their outer PBM, while 9 (69.2 %) symbiosomes contained internal vesicles. In 4 (30.8 %) symbiosomes, vesicles were attached to the outer bacteroid membrane. In summary, our integrated 2D and 3D image analyses highlight the prevalence of various vesicular structures within a continuous membrane system of symbiosomes.

**Fig. 5.**
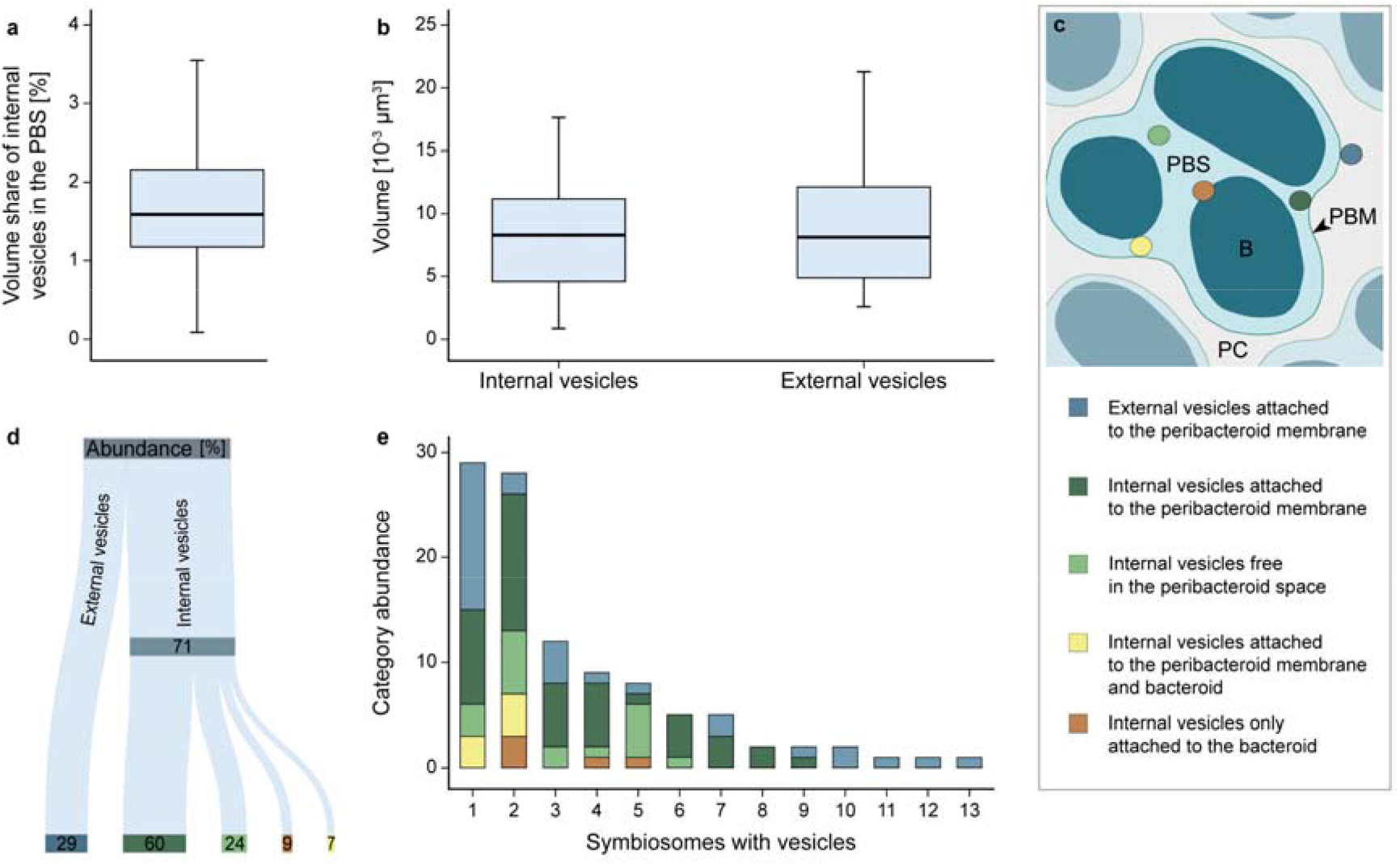
Symbiosome-related vesicle volume and abundance. **a)** PBS volume share of internal symbiosome vesicles (n = 9). **b)** Volumes of single internal (n = 75) and external (n = 30) symbiosome vesicles (Mann–Whitney U test; p = 0.406). **c)** Schematic drawing summarising the observed vesicle locations within the PBS and attached to the outer bacteroid membrane and PBM. **d)** Vesicle abundance dependent on their location inside and outside the symbiosome. **e)** Vesicle abundance by location category for each symbiosome (n = 13), ordered by descending total vesicle count.

### Membrane protrusions and vesicles with bacteroid origin

The 3D model analysis revealed vesicles inside the symbiosome compartment and in the host cytoplasm. In addition, we observed vesicle-like protrusions directly connected to the outer bacteroid membranes (Fig. 6a) and to the PBM (Fig. 6b). In some cases, these protrusions formed chain-like connections between symbiosomes, which contained small vesicles (Fig. 6c). Their diverse morphology and localisation at the symbiosome interface suggest distinct biogenesis pathways, cargo compositions, and origins, whether plant- or microbe-derived. The microbial origin of the vesicles was assessed using TEM and immunogold labelling of lipid A (LA) of the lipopolysaccharide (LPS), with 10 nm colloidal gold particles. LA is a structural component of the LPS of the outer membrane of Gram-negative bacteria, and has previously been detected in bacteroids in pea nodules (VandenBosch *et al*., 1989). Gold particles bound to LA were observed in the bacteroid cytoplasm, which is the place of LA biosynthesis (Sperandeo *et al*., 2009), at the bacteroid membranes (Fig. 6d) and the membranes of vesicles located inside (Fig. 6e) and outside (Fig. 6f) the symbiosomes. The secondary antibody did not exhibit non-specific binding. The detection of the LA signal in some of the symbiosome-associated vesicles supports that these vesicles are of microbial origin and are located at the plant-microbe interface.

**Fig. 6.**
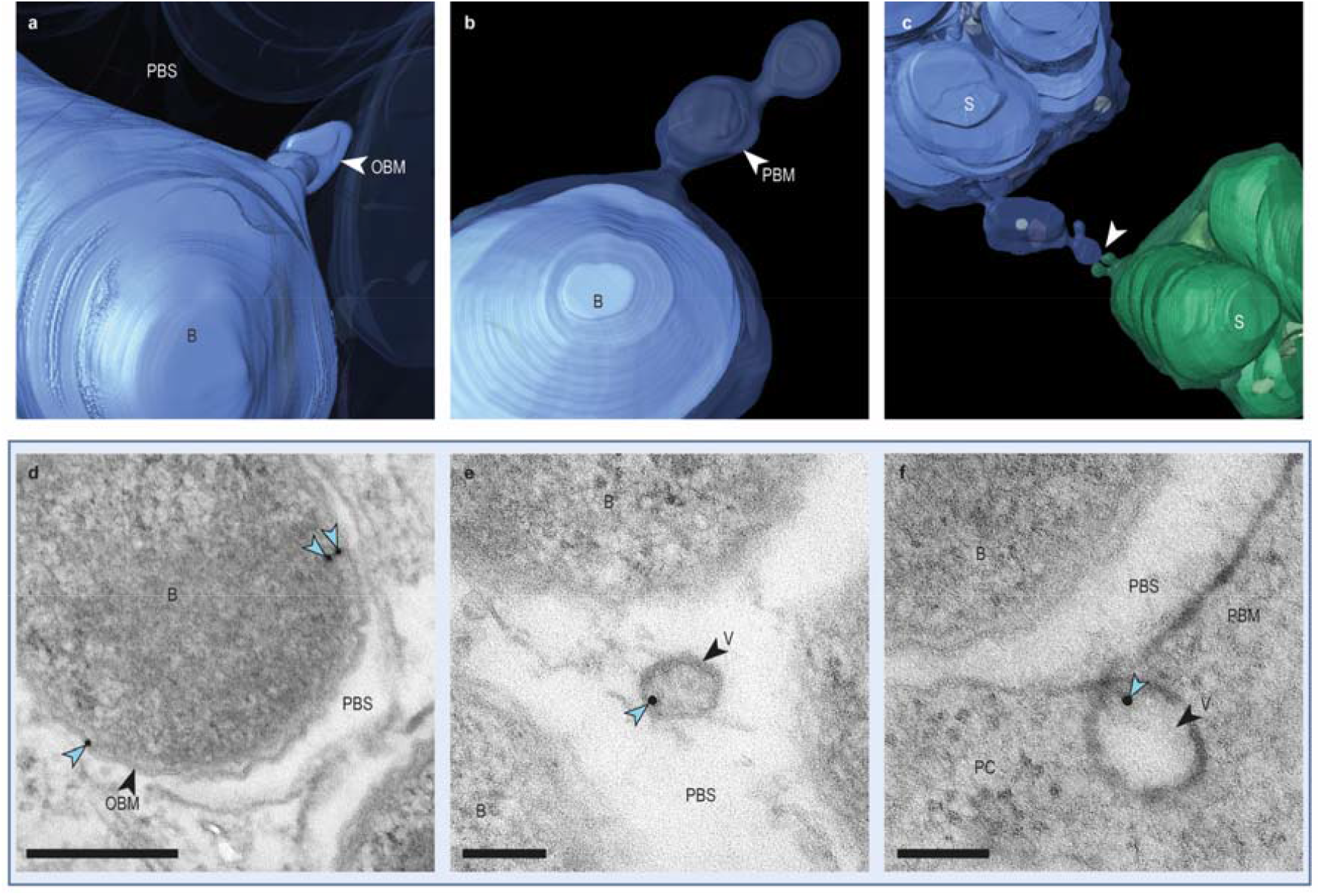
Symbiosome-associated vesicle budding and bacterial vesicle origin. **a-b)** 3D model of protruding vesicles, depicting budding or fusing events with **a)** the outer bacteroid membrane and **b)** the PBM of a symbiosome. **c)** 3D model of two symbiosomes (blue and green) with connective vesicle-like chains (white arrow). **d-f)** TEM images of symbiosome cross sections with immunogold labelling for bacterial lipopolysaccharide Lipid A. **d)** Bacteroid cross-section with gold particles (blue arrow) labelling along the membrane (scale bar: 250 nm). **e)** Cross-section of a symbiosome showing an internal vesicle labelled with a gold particle (blue arrow), located in the PBS centre (scale bar: 100 nm). **f)** Cross-section of a symbiosome showing an external vesicle labelled with a gold particle (blue arrow), attached to the PBM (scale bar: 100 nm). Labelling: B: bacteroid; PBM: peribacteroid membrane; PBS: peribacteroid space; PC: plant cytoplasm; OBM: outer bacteroid membrane, S: symbiosome; V: vesicle.

## Discussion

The intracellular accommodation of rhizobia in nodule cortical cells relies on an endocytic process in which bacteria are continuously enclosed within a plant-derived membrane compartment, a symbiosome (Tsyganova *et al*., 2021). 2D and 3D electron microscopy techniques were used to visualise *M. loti* symbiosomes formed in the central tissue of *L. japonicus* root nodules to dissect the spatial and morphological characteristics of this plant-microbe interface. The results demonstrate that the accommodation of the microbe in a symbiosome compartment did not affect the cellular rod shape of the bacterium compared to its free-living state, consistent with a previous report on determinate root nodules (Mergaert *et al*., 2006). However, morphological changes exist, as bacterial cell appendages like flagella and pili were not detected in the bacteroid state. This may reflect a reduced necessity for chemotaxis and motility once the bacteria are accommodated within a plant cell, and is supported by the observed downregulation of genes involved in chemotaxis, motility, and flagellin formation (Chen *et al*., 2023).

The accommodation of *M. loti* in symbiosomes is associated with host cell rearrangements to create space for the new cellular compartments (Fedorova, 2023). The spatial distribution of the symbiosomes is thereby organised by actin filament arrays located in the plant cytoplasm (Whitehead *et al*., 1998). The 3D reconstruction demonstrated that bacteroid orientation within the symbiosome and compartment arrangement in the plant cytoplasm are determined by the available space between existing symbiosomes and plant organelles. This likely results in their interlocking positioning. This effective occupation of space in the plant cell leads to a tight packing of small plant organelles in interstitial cytoplasmic spaces between the symbiosomes and is reflected in morphological adaptations of the organelles to the shape of surrounding compartments. It is known, that along with the symbiosis development, ER adapts its size and shape spatiotemporally (Ren *et al*., 2025). The ER, as well as dictyosomes, were frequently observed in the 3D model. This indicates sustained membrane trafficking activity in colonised cells and continuous vesicular transport to the symbiosomes (Whitehead *et al*., 1998), which persists despite severe spatial constraints in mature colonised root nodule cells because of the tight symbiosome packing. As symbiosomes with a single bacteroid were the most abundant in 2D and 3D data, this may reflect that during the symbiosome formation, bacteria are, in most cases, enveloped as single entities. This would facilitate an increase in the interface and contact area between the host and the microbe. The presence of a single bacteroid per symbiosome is a characteristic feature of indeterminate nodules, e. g. in *Medicago* (Den Herder *et al*., 2008), providing efficient host control of nutrient allocation to individual bacteroids. Symbiosomes containing multiple bacteroids may result either from a simultaneous release in a symbiosome compartment or from bacterial cell division occurring within symbiosomes (Goodchild & Bergersen, 1966), as demonstrated by TEM. Despite the large sample size of the 2D dataset, the number of symbiosomes containing multiple bacteroids can be underestimated due to the limited morphological information provided by a single focal plane (Harwood *et al*., 2020). However, multi-bacteroid symbiosomes may be able to convert into compartments with a single bacteroid, as the tubular connections formed by the PBM may represent residual structures from symbiosome division. The PBM contours the shape of the microbe and is often in direct contact with it. Therefore, the shape and size of a symbiosome highly depend on the bacteroid orientation and the total bacteroid volume it includes (Studer *et al*., 1992). The PBS is present only as a thin layer between the bacteroid and the PBM and occupies a small volume of the symbiosome compartment. It is known that the PBS contains substances of host and microbial origin and that transport and signalling between the two partners must occur through this matrix (Ayala-Garcia *et al*., 2025). Therefore, it may be advantageous that the PBS layer is relatively thin, minimising the distance between the microbe and the plant once the selectively permeable, physical PBM barrier has been crossed.

The documentation of chain-like evaginations and tubular connective structures formed by the PBM may hint towards a dynamic and flexible system that allows cross-linkage and exchange mechanisms among independent symbiosomes and their PBS. Consequently, inter-entity transfer could be possible. With regard to inter-organismal plant-microbe communication and nutrient transfer, an exchange of substances across PBS and PBM appears necessary to enable a successful symbiosis (Reagan *et al*., 2018). Vesicles with heterogeneous cargo, documented inside and outside the symbiosome compartment, may carry a wide spectrum of substances and proteins (Ayala-Garcia *et al*., 2025). They can be part of the transport system between the two organisms as an extracellular vesicle-driven crosstalk through the PBS (Ayala-Garcia *et al*., 2025). Extracellular vesicle-mediated transport is one of the major pathways for the cross-kingdom interaction, as they are considered as effective carriers, e.g. of biologically active components such as proteins, lipids and nucleic acids for intercellular communication in prokaryotes and eukaryotes (Liu *et al*., 2021). Furthermore, outer membrane vesicles derived from gram-negative bacteria can play a major role in plant immunity modulation and bacterial cell-cell communication (Katsir & Bahar, 2017).

The identification of bacterial LA as a component of vesicular membranes provides evidence that some vesicles within and outside the symbiosome have a bacteroid origin. This evidence is reinforced by the documentation of membrane evaginations along the outer bacteroid membrane, which may represent a vesicular budding event. How bacterial vesicles reach the plant cytoplasm and are transported across the PBM barrier remains to be explained. Since in the TEM images, gold particles were rarely present along the PBM, it is unlikely that vesicles with bacteroid origin fuse with the PBM and release content into the plant cytoplasm. However, with regard to the abundance of MVBs between the symbiosomes and the documentation of MVB-like compartments connected to symbiosomes in the 3D model, it seems possible that bacterial vesicles can bud off the symbiosome corpus and are transferred to the plant cytoplasm.

There is no experimental evidence that unlabelled vesicles have a plant origin, and the absence of gold labelling on certain vesicles may be attributed either to a genuine lack of LA or to limited epitope accessibility at the surface of the ultrathin section during sample preparation (Hyatt & Wise, 2001). However, MVB of plant origin could fuse with the PBM. Similar events have been observed at the plant-fungi interface in arbuscular mycorrhiza symbiosis (Roth *et al*., 2019). Furthermore, the documented external vesicles attached to the PBM from the plant cytoplasm side and the vesicle-like PBM evaginations can hint at fusion events of vesicles with plant origin that open towards the PBS (Kijne & Pluvqué, 1979). As PBS is known to include substances from plant origin, this could be a possible option for the plant to deliver compounds to the accommodated microbe apart from transporters and channels in the PBM (Jeong *et al*., 2004; Clarke *et al*., 2014). Intense vesicle trafficking in the plant cell helps deliver cargo such as proteins or secondary metabolites directly to the interaction site (Bapaume & Reinhardt, 2012). Furthermore, vesicle fusion may contribute to the membrane formation of the symbiosome envelope (Roth & Stacey, 1989; Fedorova, 2023). Although the question whether vesicular structures are fusing with or budding off the PBM cannot be answered in this study, the 2D and 3D data enlightened that membrane modifications especially of the PBM interface are common for this symbiosis and may play an important role in the plant-microbe interaction and communication (Kijne & Pluvqué, 1979; Ayala-Garcia *et al*., 2025).

In summary, this study provides an ultrastructural analysis of the arrangement and morphology of symbiosomes and their related vesicular structures in *L. japonicus* root nodule symbiosis, by combining TEM and FIB/SEM as complementary techniques. This study expands on previous research on the ultrastructure of arbuscules in cortical cells of host roots (Ivanov *et al*., 2019; Roth *et al*., 2019), offering a comparative view of the symbiotic plant-microbe interface. The reconstruction of symbiosome interfaces in 3D, delivered a multi-angled view of specific structures, enabled to determine the volumes, spatial distribution, and connectivity of symbiosomes and their associated features that cannot be fully addressed with 2D image data. However, 2D TEM images are a valuable approach for estimating symbiosome structural proportions and origins based on area measurements and well-established labelling techniques. Future research with focus on the symbiosome interface across diverse microbe-host combinations could help to unravel mechanisms and structural features facilitating the efficiency of this symbiosis. In addition, a deeper understanding of the vesicle-based interplay between the host and rhizobia can yield new insights into the maintenance of this complex interaction. Ultimately, ultrastructural dissecting of this symbiotic dialogue provides an opportunity to assess new strategies for sustainable agriculture (Goyal *et al*., 2021).

## Acknowledgements

We thank the Transregio TRR 356 of the German Research Council (DFG) and the colleagues of the electron microscopy facility at the biology faculty LMU Munich for supporting this project. Furthermore, we thank Kareem Farghaly, who contributed to the 2D TEM data analysis of symbiosomes.

## Competing interest

The authors have declared that no competing interests exist.

## Author contributions

IG coordinated the project, did the sample preparation and microscopy, wrote the manuscript and created all figures and drawings. KP contributed to the sample preparation and the text. AK coordinated the project and contributed to the acquisition of the FIB/SEM data set.

## Funding

This work was supported by the Transregio TRR 356 (Project number 491090170) of the German Research Council (DFG) with TP-B07 to KP and TP-Z02 to AK. The funder had no role in study design, data collection and analysis, decision to publish, or preparation of the manuscript.

## Notes

### Competing Interest Statement

The authors have declared no competing interest.

