## Supplementary figures for "The 2D and 3D ultrastructure of symbiosomes and associated vesicular structures in *Lotus japonicus* root nodule symbiosis"

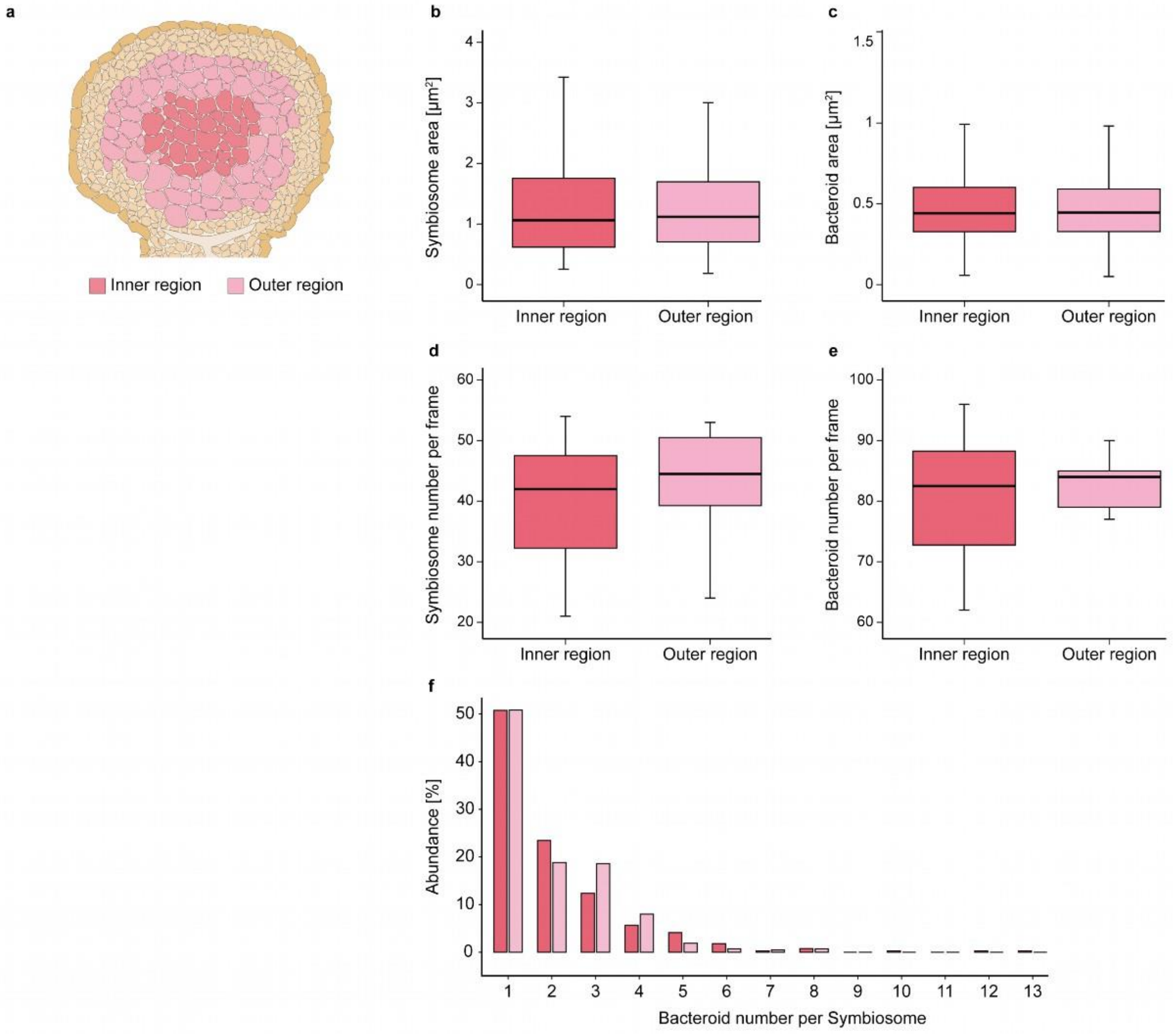


**Supplementary Fig. 1 | Comparison of symbiosome characteristics between inner and outer regions of colonised *L. japonicus* root nodule.** The graphs represent quantitative data derived from TEM images of the inner and outer regions (ten cells each) of a colonised root nodule tissue of a single root nodule. **a)** Schematic drawing of a root nodule cross-section with inner and outer colonised regions highlighted in different colours. **b)** Symbiosome area in cells from the inner (n = 390) and outer (n = 428) tissue regions (Mann–Whitney U test; p = 0.316). **c)** Area of individual bacteroid cells in the inner (n = 806) and the outer (n = 862) tissue region (Mann–Whitney U test; p = 0.700). **d)** Number of symbiosomes in the inner and outer region per image frame (n = 10) (t-Test; p = 0.443). **e)** Number of bacteroids in the inner and outer region per image frame (n = 10) (t-Test; p = 0.296). **f)** The abundance of symbiosomes with a given bacteroid number (from 1 to 13) in the inner (n = 390) and outer (n = 428) tissue region.

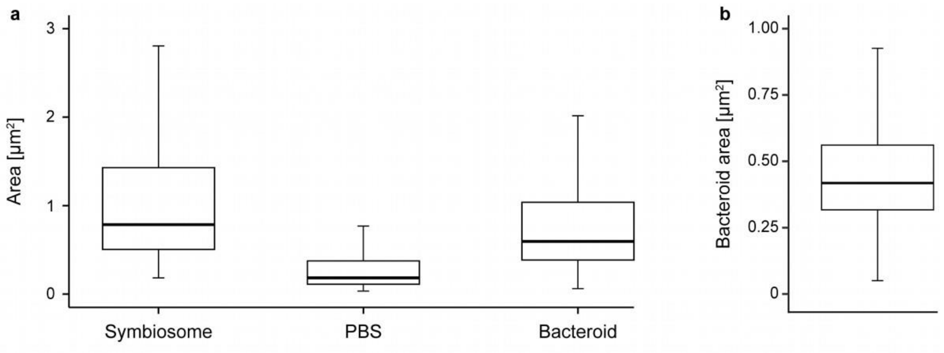


**Supplementary Fig. 2 | Area of symbiosome-related structures in the outer colonised tissue layer in *L. japonicus* root nodules.** Measurements generated by TEM images of the outer colonised cell region of three root nodules. **a)** Area distribution of symbiosomes (n = 1512), their PBS and their total bacteroid area. **b)** Area of single bacteroid cells (n = 2584).
